# A multidisciplinary approach to a unique Palaeolithic human ichnological record from Italy (Bàsura Cave)

**DOI:** 10.1101/529404

**Authors:** Marco Romano, Paolo Citton, Isabella Salvador, Daniele Arobba, Ivano Rellini, Marco Firpo, Fabio Negrino, Marta Zunino, Elisabetta Starnini, Marco Avanzini

## Abstract

Based on the integration of laser scan, sedimentology, geochemistry, archeobotany, geometric morphometrics and photogrammetry, here we present evidence testifying a Palaeolithic group that explored a deep cave in northern Italy about 14 ky cal. BP. Ichnological data enable us to shed light on individual and group level behavior, social relationship and mode of exploration of the highly uneven environment. Five individuals, two adults, an adolescent and two children, entered the cave barefoot and with a set of wood chips to illuminate the way. Traces of crawling locomotion are documented for the first time in the global human ichnological record. The anatomic details recognizable in the crawling traces show that no garment was interposed between the limb and the trampled sediments. Our study demonstrate that very young children (the youngest less than three years old) were active members in the Upper Palaeolithic populations, even in seemingly dangerous activities and social ones.

## Introduction

Discovered in 1950, the ‘Grotta della Bàsura’ is a large cave that has produced some of the most important Italian prehistoric human research discoveries of the twentieth century (Chiappella, 1953; Tongiorgi and Lamboglia, 1954; Blanc, 1960; Lamboglia, 1960; Giacobini, 2008). The cave opens at 186 m a.s.l. about 1 km north of Toirano (Savona, Italy - 436253.433 E; 4887689.739 N), and develops in the core of Mount S. Pietro for 890 linear meters with a height difference of + 20 / - 22 meters with respect to the entrance. Originally, abundant remains of *Ursus spelaeus* and numerous traces of human attendance (footsteps, charcoals, digit tracks, lumps of clay adhering to the walls) were found in different areas of the cave within approximately 350 m of the entrance. Regrettably, destruction of most of the ichnological record occurred due to uncontrolled cave visits following discovery (Blanc et al., 1960; De Lumley and Giacobini, 1950).

The first scientific study of the ‘Grotta della Bàsura’ concerned the analysis of thirteen footprints attributed on morphometric basis to a Neanderthal population (Pales, 1960). The association, in the last room of the cave called *‘Sala dei Mister’* (Mystery’s room), between human footprints and finger flutings on a clay-coated stalagmite plus lumps of clay adhering to the cave walls, as well as a consistent deposit of cave bear bone called *‘Cimitero degli Orsi’* (Cave-bear’s cemetery) (Chiappella, 1952) lead workers to hypothesize that Neanderthals attended the cave for hunting cave bears and ritual purposes (Blanc, 1960).

Radiometric dating on charcoal samples from the trampling palaeosurface constrains the exploration of the cave in a limited time span of the late Upper Palaeolithic, between 12,310±60 and 12,370±60 (Table 1).

**Table 1.** Radiometric dating of charcoals collected from the trampling palaeosurface during 2017 excavations inside the ‘Sala dei misteri’. (*14C ages have been calibrated to calendar years with software program: OxCal, version 4.3. Used calibration curve: IntCal13)

| SAMPLE NAME | PROVENANCE | DATED MATERIAL | DESCRIPTION | LAB. CODE | F<sup>14</sup>C ± 1σ | <sup>14</sup>C AGE (yr BP)<br>± 1σ | CALIBRATED*<br>DATING RESULT<br>(95.4 probability) | %C | %N | δ<sup>13</sup>C<br>(in ‰; ±1σ) | δ<sup>15</sup>N<br>(in ‰; ± 1σ) | C/N ratio |
| --- | --- | --- | --- | --- | --- | --- | --- | --- | --- | --- | --- | --- |
| Bàsura SM B5 17B | ‘Sala dei misteri’, square B5, Unit 1 | charcoal (AAA) | *Pinus t. sylvestris/mugo* | GrA-69598 | 0.2160±0.0016 | 12310±60 | 12720–12110 cal BC | 71.4 | - | -<br>25.24±0.14 | - | - |
| Bàsura SM D6 33B | ‘Sala dei misteri’, square D6, Unit 1 | charcoal (AAA) | *Pinus t. sylvestris/mugo* | GrA-69597 | 0.2145±0.0016 | 12370±60 | 12830–12165 cal BC | 63.3 | - | -<br>26.37±0.14 | - | - |

Since 2014, a multidisciplinary study was promoted under the direction of two of the present authors (ES, MZ). The aim of this work was to review and improve the knowledge of the Grotta della Bàsura’s human ichnology, soil micromorphology, sedimentology and radiocarbon chronology. A preliminary report from this study focused on the human footprints preserved in the innermost room of the cave (Citton et al., 2017). In the present paper, we broaden our study to analyze and interpret all human traces *sensu lato* from the ‘Grotta della Bàsura’, providing new insights about the behaviour, the identity of the members, the exploratory techniques, and the social structure of an Upper Palaeolithic group.

### Data

A total of 179 footprints and traces *sensu lato* were recorded and considered (Table 1). In addition to footprints (Figure 1), among the traces were unintentional digit and hand traces on the clay-rich floor and smears from hands dirtied with charcoal on the side walls of the cave. Charcoal remains with bundles of *Pinus sylvestris/mugo* originally utilized with illuminated the cave are preserved on the trampling palaeosurface and on the floor of the *‘Sala dei Mister’*.

**Fig. 1.**
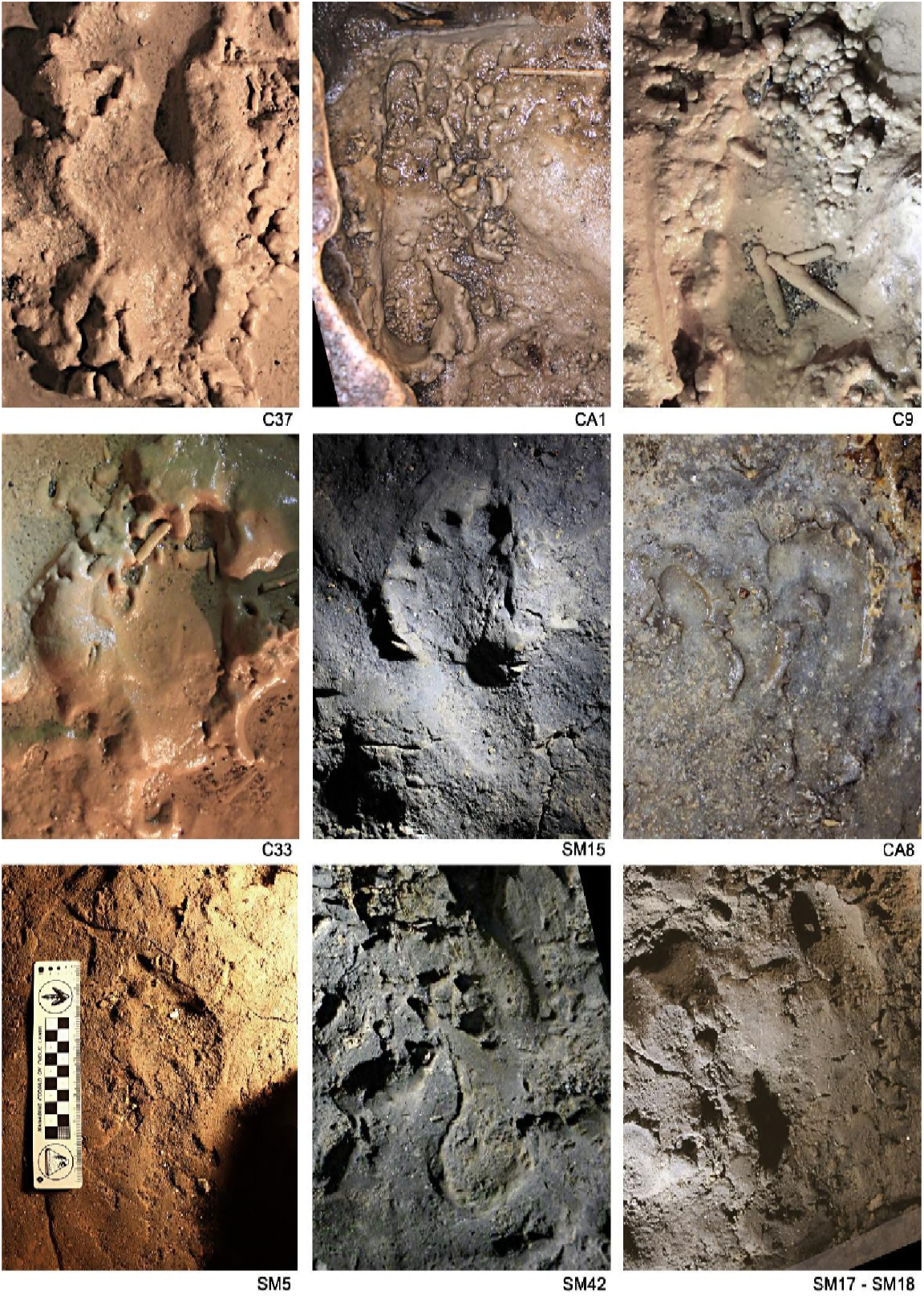
Human footprints imprinted on muddy substrate in different moisture conditions. C37, Human footprint referred to the Morph 5 (‘lower corridor’). CA1 and C9, Human footprint referred to the Morph 4 (‘upper corridor’). C33, Human footprint referred to the Morph 3 (‘lower corridor’). SM15, Human footprint referred to the Morph 3 (‘Sala dei Misteri’). CA8, Human footprint referred to the Morph 3 (‘upper corridor’). SM5 and SM42, Human footprint referred to the Morph 2 (‘Sala dei Misteri’). SM17 and SM18, Human footprint referred to the Morph 1 (‘Sala dei Misteri’).

Other digit and hand traces indicating clay excavation and finger flutings (e.g. SM53, SM44; Figure 2), probably related with social or symbolic activities, can be instead considered intentional. Moreover, different sized bear and Canidae *incertae sedis* footprints are ubiquitously present, and are often associated with human prints.*Ursus* sp. hibernation areas are still recognizable with well-preserved nests of both cubs and adult bears.

**Fig. 2.**
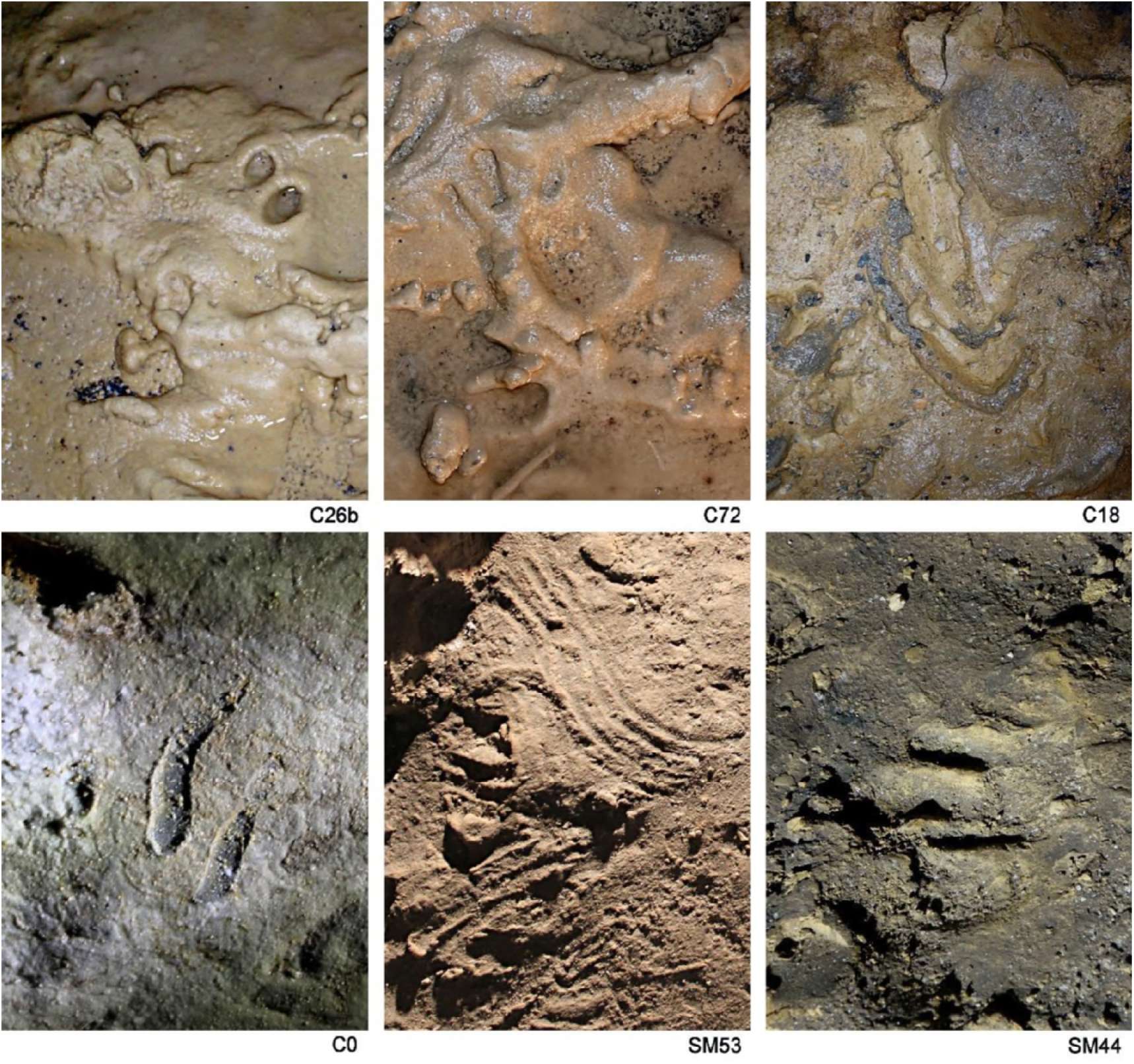
Finger and hand prints on muddy substrates. C26b, Finger traces (‘lower corridor’). C72, Hand print (‘lower corridor’). C18, Undetermined trace (‘lower corridor’). C0, Two finger traces on the concretioned side-wall of the ‘lower corridor’. SM53, Finger flutings on a clay-coated stalagmite (‘Sala dei Misteri’). SM44, finger traces (‘Sala dei Misteri’).

### Preservation

Footprints are preserved in several areas of the cave, particularly in the innermost room (*‘Sala dei Misteri’*) and in the main gallery (*‘Corridoio delle impronte’*), which is divided into two corridors at different topographic heights of about 5 m (hereafter, lower and upper corridor respectively) (Figure 3).

**Fig. 3.**
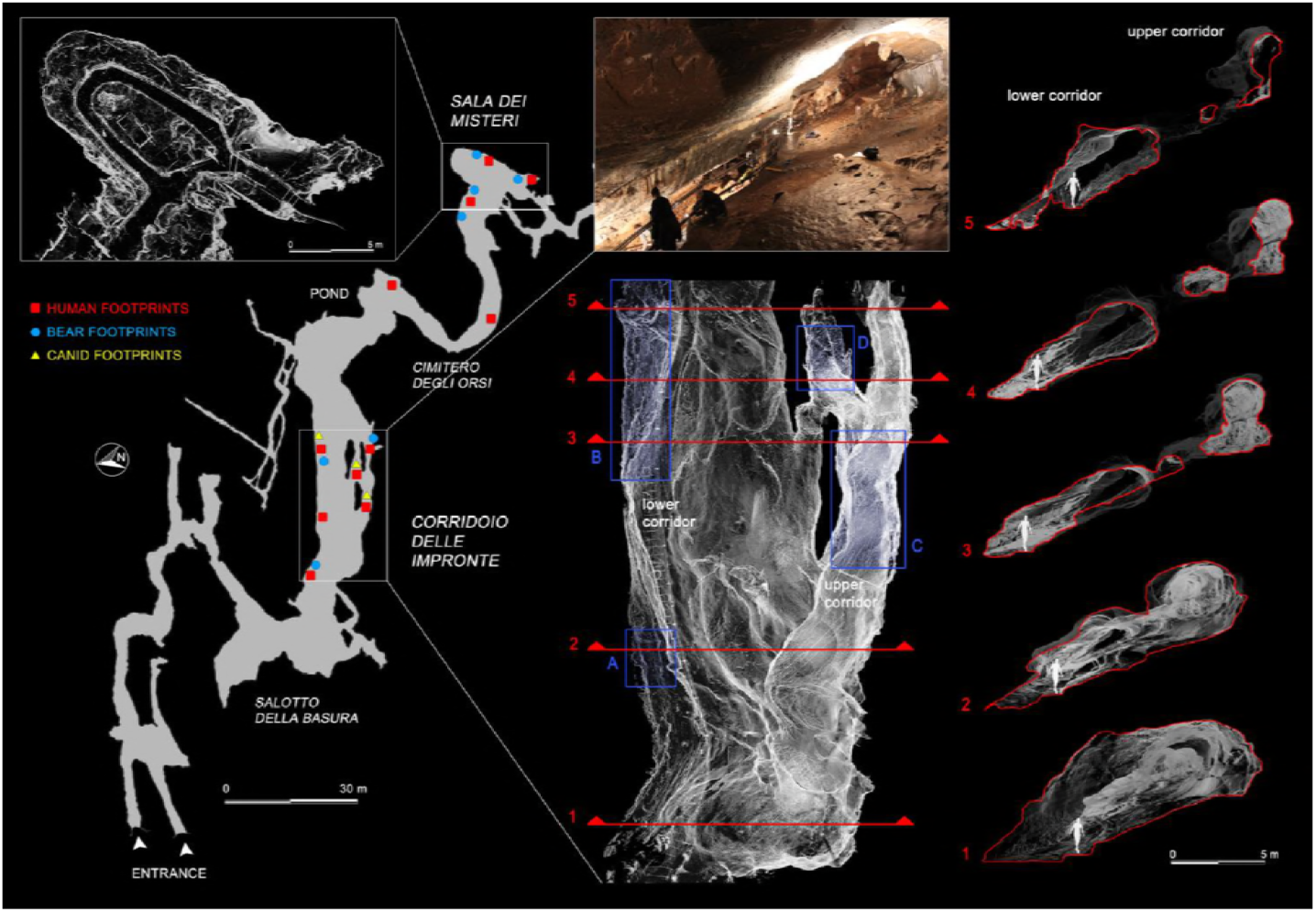
Planimetry of the *‘Grotta della Bàsura’* and location of human, bear and canid footprints. White rectangles enclose the three-dimensional reconstructions, obtained via laser scanner, of the innermost room (*‘Sala dei Misteri’* - left) and the main gallery *(‘Corridoio delle impronte’* - right) of the cave, where the human footprints are preserved. Cross-sections obtained from the three-dimensional reconstruction of the main gallery are highlighted in red and show the branching of the ‘lower’ and ‘upper’ corridors, respectively. Blue rectangle indicate the four areas within the main gallery where most of the human footprints are concentrated (A and B for the lower corridor, C and D for the upper corridor).

The flooding dynamics and cave geometry produced two contrasting situations for sediment deposition and transport inside the cave. Detrital sediments, constituted by silty clay and well-sorted sandy sediments, are most abundant on the floor of the *‘Sala dei Misteri’*. The coarse elements, defined as grains of gravel size and larger (>2 mm), consist mainly of few biogenic components (fragments of bear bones). The sandy fraction is comprised of allogenic, surface-derived siliciclastic sediments (alluvium). The *‘Sala dei Misteri’* appear to undergo episodic filling and excavation controlled by catastrophic storms.

The sediments in the *‘Corridoio delle impronte’* show a high proportion of mud with many coarse lithic fragments and are mainly carbonates (calcite and dolomite), suggesting an autogenic origin. Here, the trampled substrate is poorly consolidated and superimposed on a stalagmite crust. At the time when humans and other large mammals left their impressions, different conditions of the substrate co-existed even in small sectors of the cave. In some areas, the substrate was in a highly plastic condition, whereas in other it was waterlogged or submerged. Different water contents account for the variable detailing of tracks (e.g. registration of plantar arch, heel and metatarsal regions, digit tips, track walls) and in particular of the associated extra-morphologies (e.g. expulsion rims, slipping traces). The surface and the footprints are cross-cut by mud cracks, suggesting a loss of moisture in sediments after trampling. Carbonate crusts (composed of both calcite and dolomite) cover many of the footprints in areas subject to more intense dripping. Iron and manganese oxide coatings were found in the crust, due to repeated immersion in ponded water.

## Results

Geometric morphometry performed on human footprints highlighted five main morphotypes (hereafter Morphs.) indicating a possible minimum number of five individuals attending the cave (Figure 4). This number is confirmed by the construction of morphological groups (Table 2) reconstructed through the overlapping of footprints that show a variability less than 2% of the main parameters.

**Fig 4.**
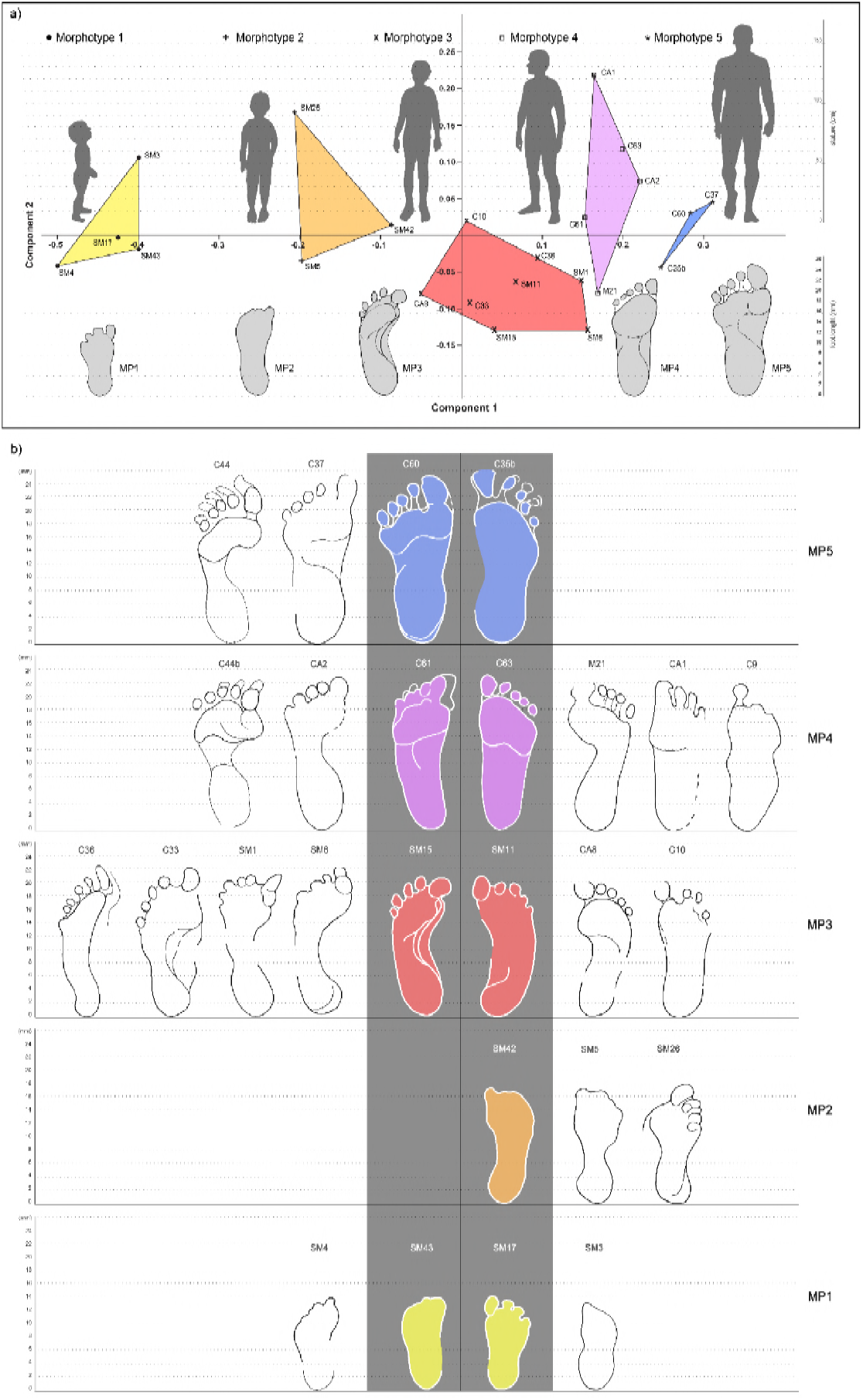
Principal Component Analysis based on the best-preserved footprints from *‘Sala dei Misteri’* and *‘Corridoio delle impronte’*. **a**, The five morphotypes to which footprints have been referred are shown above. **b**, Selected outlines of the best preserved footprints, for each recognized morphotype, are reported.

**Table 2.** Footprint shape features after “Robbins footprint recording form” (1985, p.97-102).

| ID | Left (L) or right (R) foot | General appearance |  | Relative length of toes | Toes region general appearance length - width | Toe 1 position | Ball region, general appearance length-width | Arch region |  | Heel region, general appearance | Heel posterior margin |  |
| --- | --- | --- | --- | --- | --- | --- | --- | --- | --- | --- | --- | --- |
|  |  | Length | Width |  |  |  |  | Medial margin | Lateral margin |  |  |  |
| SM3 | R | short | broad | 1, 2, ? | short - broad | extended, anteriorly |  | straight | concave | circular | convex pronounced | Morphotype 1 |
| SM4 | L |  |  | 1,2,3,4,5 | short - broad | extended, anteriorly | short - moderate |  | concave |  |  |  |
| SM43 | L | short | broad | 1,2, ? | short - broad | extended, anteriorly |  | straight | convex |  | convex pronounced |  |
| SM17 | R | short | broad | 1,2,3,4,5 | short - broad | extended, anteriorly |  | straight/concave | convex |  | convex pronounced |  |
| SM5 | R | moderate | moderate | 1,2,3,4,5 | short - broad | extended oblique medially | long - moderate | concave | concave |  | convex slight | Morphotype 2 |
| SM42 | R | moderate |  | 1,2, ? |  | extended oblique medially |  | concave |  |  | convex pronounced |  |
| SM26 | L | moderate | broad | 1,2,3,4,5 | short - moderate | extended oblique laterally |  | concave | concave |  | convex slight |  |
| CA8 | R | long | moderate | 1,2,3,4,5 | moderate - broad | extended oblique medially | long - moderate | concave | straight | oblong | convex pronounced | Morphotype 3 |
| CA10 | R | long | moderate | 1,2,3,4 | moderate - broad | extended oblique medially |  | straight/concave | straight | oblong | convex pronounced |  |
| SM15 | L | long | moderate | 2,3,1,4,5 | moderate - broad | extended oblique medially | long - narrow | concave | straight | oblong | convex pronounced |  |
| SM11 | R | long | moderate | 1,2,3,4,5 | short - broad | extended anteriorly |  | straight/concave | convex | oblong | convex pronounced |  |
| SM6 | L | long | moderate | 2,1,3,4,5 | short - broad | flexed slight | long - narrow | concave | convex | oblong | convex pronounced |  |
| SM1 | L | long | moderate | 2,3,1,4,5 | short - broad | extended oblique medially | moderate - narrow | unknown | convex | circular | convex pronounced |  |
| C33 | L | long | broad | 1,2,3,4,5 | long - broad | extended oblique medially | long - broad | concave | convex | oblong | convex pronounced |  |
| C36 | L | long | very narrow | 1,2,3,4,5 | moderate - narrow | extended oblique laterally | long - narrow | concave | straight | oblong | convex pronounced |  |
| CA1 | R | long |  | 1,2,3,4 | moderate - broad | extended anteriorly | moderate - narrow | straight |  | oblong | convex, pronounced | Morphotype 4 |
| CA2 | L | long | moderate | 1,2,3,4,5 | moderate - broad | extended anteriorly | moderate - narrow | concave | straight | oblong | convex pronounced |  |
| C61 | L | long | moderate | 1,2,3,4,5 | moderate - broad | extended oblique medially | moderate - narrow | straight/concave | straight/convex | oblong | convex pronounced |  |
| C63 | R | long | moderate | 1,2,3,4,5 | moderate - broad | extended anteriorly | moderate - narrow | straight/concave | straight/concave | oblong | convex pronounced |  |
| M21 | R | long | moderate | 1,2,3,4,5 | moderate - broad | extended anteriorly | moderate - narrow | concave | straight | oblong | convex pronounced |  |
| C9 | R | long | moderate | 1 |  | extended anteriorly | moderate - narrow | straight | convex | oblong | convex pronounced |  |
| C44b | L | long | broad | 2,1,3,4,5 | short - broad | extended oblique medially | moderate - broad | concave | convex | oblong | convex, moderate |  |
| C60 | L | long | broad | 1,2,3,4,5 | moderate - broad | extended oblique laterally | moderate - broad | straight/concave | straight/concave | circular | convex moderate | Morphotype 5 |
| C37 | L | long | broad | 1,2,3,4,5 | moderate - broad | extended anteriorly | moderate - broad | straight/concave | straight/concave | circular | convex moderate |  |
| C35b | R | long | broad | 1,2,3,4,5 | moderate - broad | extended anteriorly | moderate - broad |  | straight/concave | circular | convex moderate |  |
| C44 | L | long | broad | 1,2,3,4,5 | moderate - broad | extended anteriorly | moderate - broad | concave | straight/concave | oblong | convex pronounced |  |

Morphs. 1 and 2 can be easily distinguished on the basis of the absolute footprint size. Morph. 1 includes footprints with a length of 13.55±0.49 cm showing characters indicative of an early ontogenetic stage of the producer, such as digit traces and heel area proportionally wider than longer tracks. Morph. 2, with a length of 17 cm, is distinguished by Morph. 1 on the basis of a more pronounced plantar arch. Morph. 3 comprises footprints 20.83±0.51 cm in length (Figure 4 and Table 3). A plantar area characterized by a very pronounced medial embayment, to which correspond a strongly convex external margin, is shared together with a strong adduction of digit I trace, an overall larger divarication of digit traces and a consistent separation between adjacent digits II-III and IV-V.

**Table 3.**
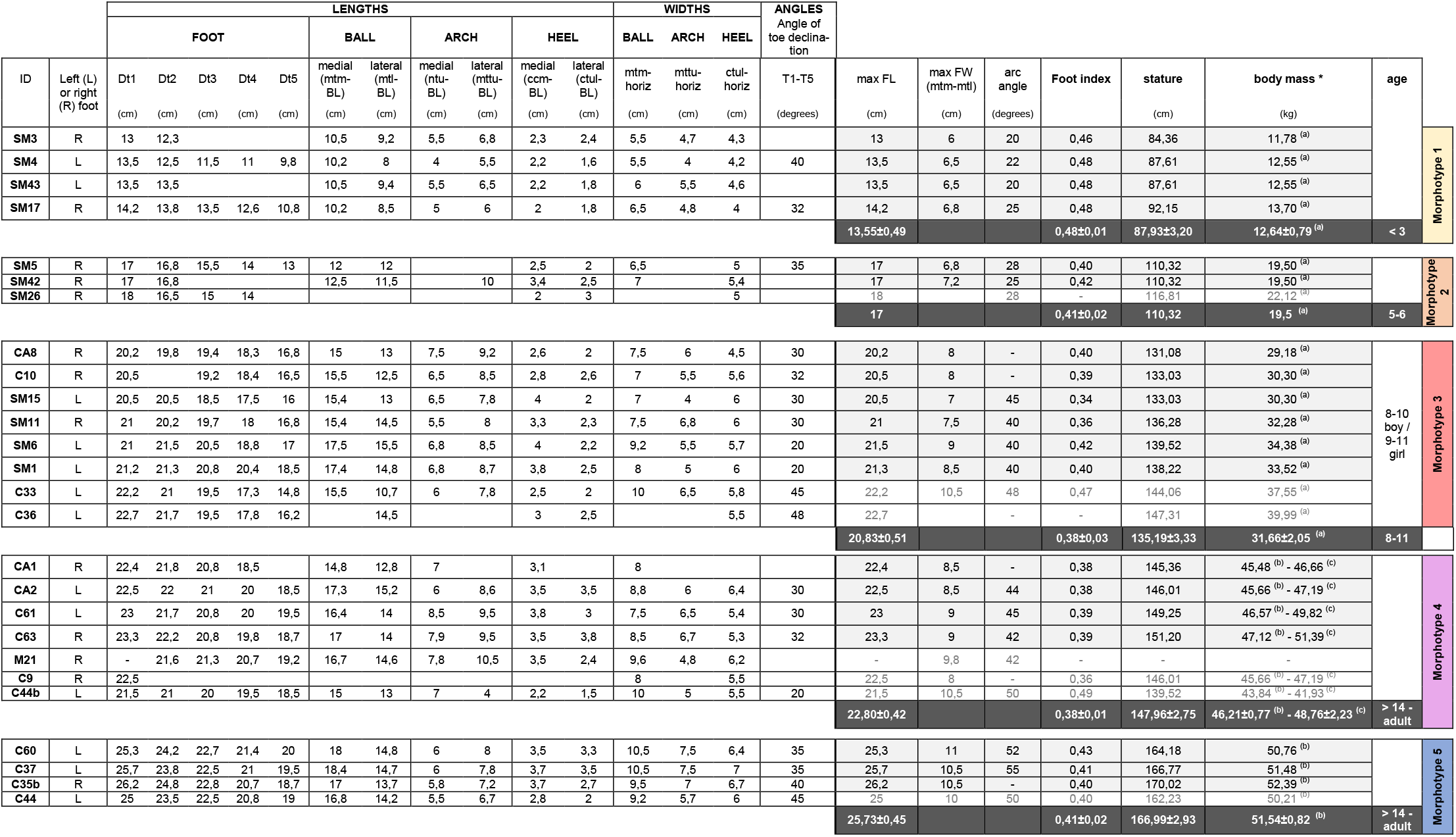
Measurements and elaboration data (foot index, stature, body mass and age) based on the best-preserved tracks from the ‘Sala dei Misteri’ and ‘Corridoio delle impronte’. *Body mass: (a) Citton et al. 2017; (b) Bavdekar et al. 2006; (c) Grivas et al. 2008 (see text).

Morph. 4 (Figures 4, 5) is represented by bigger footprints (22.80±0,42 cm in overall length) with roughly straight medial and lateral margins and a medial embayment less marked than in Morph. 3. Digit tip traces are strongly aligned and oriented forward as the footprint axis does. Morph. 5 (Figures 4, 6) includes footprints of 25.73±0.45 cm in overall length and slightly concave margins, with a variably pronounced plantar embayment. The footprints appear generally more robust and stockier with respect to Morphs. 3 and 4, sharing with Morph. 4 the straight, forwardly oriented digit tips and with Morph. 3 an adducted digit I trace.

**Fig. 5.**
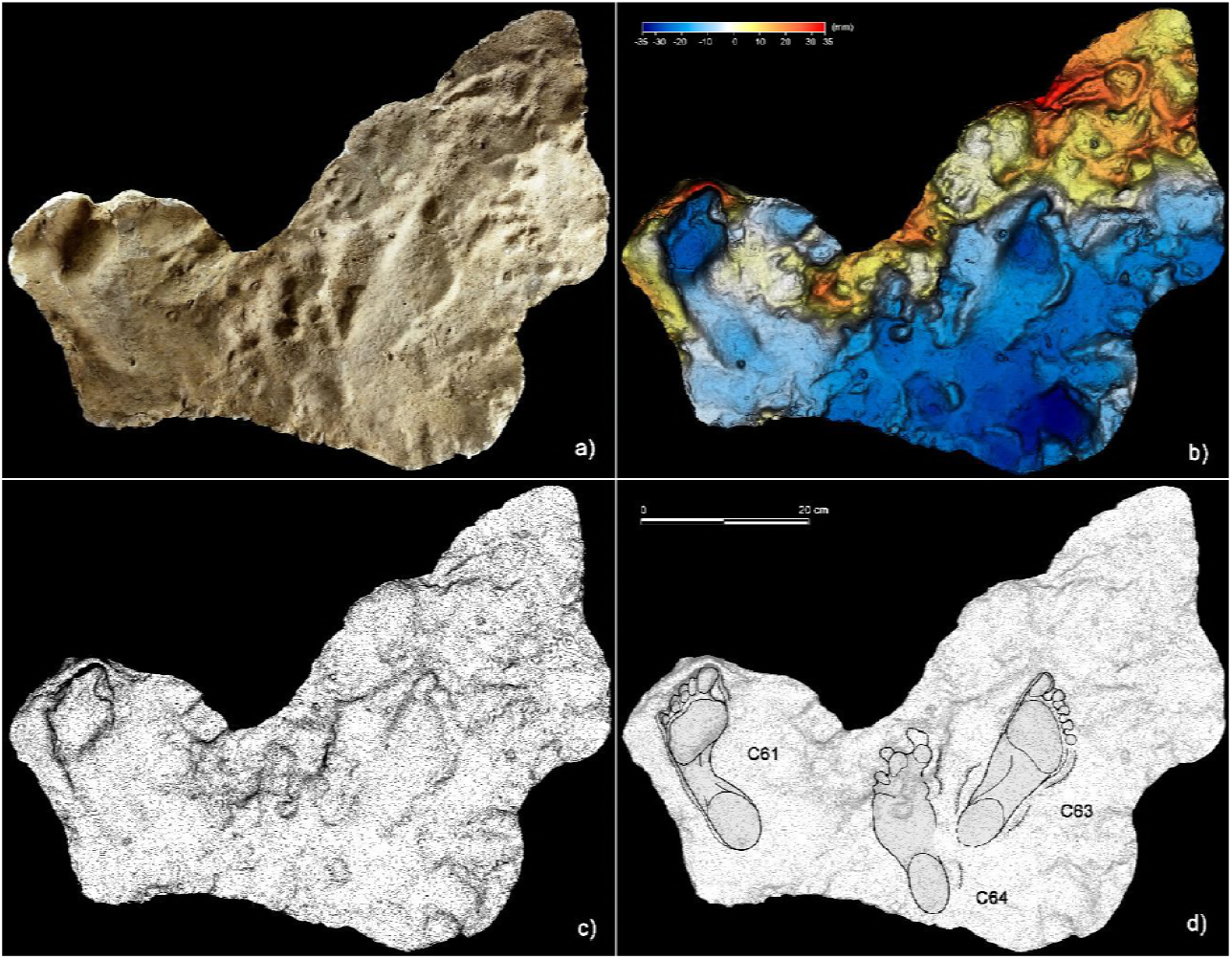
Plantigrade tracks from the ‘lower corridor’. **a**, cast of the 1950s reproducing tracks C61, C63 and C64, preserved in the sector A of the ‘lower corridor’ (see Fig. 1 main text). **b**, Digital terrain model of the cast obtained from the HDI 3D Scanner. **c**, Topographic profile with contour lines, obtained from b. **d**, Interpretive draw. Note that the tracks C61 and C63 were most likely left by a producer (Morph. 4) crouched against the side-wall of the ‘lower corridor’.

Plantigrade tracks allowed us to estimate stature, weight and ontogenetic stage of the producers basing on the collected biometric measurements (Table 3) and the adopted formulas (see Methods). An estimation of the gender for Morph. 5 was also attempted (see Methods).

The group of track producers entering the cave comprised: a three-year-old child about 88 cm tall (Morph. 1); a child at least six years old and about 110 cm tall (Morph. 2); a pre-adolescent, between eight and eleven years old, about 135 cm tall (Morph. 3); a sub-adult to adult about 148 cm tall (Morph. 4); and an adult about 167 cm tall (Morph. 5). Estimate of the stature for the Morph. 5 is also sustained by the results obtained considering the length of the tibia derived from the available kneeling traces (see Methods). Our results concerning Morphs. 4 and 5, which are referred to adult individuals, are in agreement with the average stature suggested during the European Upper Palaeolithic (162.4±4.6 cm for males and 153.9±4.3 cm for females) (Villotte et al., 2017).

Digitigrade and semi-plantigrade footprints informed on the pedal postures and the behaviour of the producers passing through different sub-environments of the cave. Both these footprint types were in most cases traced back to the same type of producer by comparison with complete footprints indicating complete foot support during locomotion. Some semi-plantigrade footprints (e.g. C44, C44b; Figure 7) show a strongly adducted trace of digit I and an apparent alignment with the other digits, probably because of the intra-rotation movements of the distal portion of the foot during the thrust phase. Footprints included in the Morph. 3 show a peculiar pedal morphology. While the resulting morphology of the digit I trace is explained by walking on a waterlogged substrate, the separation between digit pairs II-III and IV-V allowed hypothesizing an inherited familiar trait or a pathological condition of the producer’s feet. The producer was not invalidated, showing the greatest mobility in the hypogeal environment, an evidence that could refer to a ‘hyperactive’ exploratory behavior typical of the pre-adolescent/adolescent stage.

In the lower corridor (Figure 8), a few of these footprints are clearly associated with elongated traces imprinted by the producers’ knees resting on the substrate. Successions of kneeling traces can be clearly recognized for Morphs. 3, 4, and 5. Based on the overall size of the metatarsal and knee couples aligned on the substrate, crawling deambulation in a totally unknown environment is inferred for the whole group. These knee imprints (e.g. C42; Figure 7) show the completely muscular structure of the joint and of the next regions. The patella, the patellar ligament (tendon), the tibial tuberosity, the fibular head, the basis of the vastus medial and the iliotibial band are recognizable and allow us to infer the body structure of the trackmakers.

## Discussion

Successions of kneeling traces allowing one to infer a crawling locomotion for the trackmakers have never been reported before. Isolated kneeling traces (i.e. a footprint followed by a knee imprint of the same leg) have only previously been reported from the ‘Galerie Wahl’ at Fontanet cave and from the ‘Salle des Talons’ at Tuc d’Audoubert cave, France (Pastoors et al., 2015) but are not sufficient to infer crawling deambulation. In addition, the anatomic details clearly recognizable in the crawling traces from ‘Grotta della Bàsura’ enabling hypothesize that no garment was interposed between the limb and the trampled sediments.

The integration of all the available ichnological evidence in the complex morphology of the cave enabled us to reconstruct a detailed hypothesis for the events occurred during the exploration of the cave about 14,000 years ago by Paleolithic peoples. Five individuals, comprising two adults, an adolescent and two children, entered the cave barefoot and with a set of *Pinus sylvestris/mugo* bundles to be burned for illumination. The adopted lighting system would allow a longer time of lighting, as inferred from the fire wood illumination bundles adopted by the Bronze age salt miners at Hallstatt (Grabner et al., 2007, 2010). Lightning bundles are usually made of resinous wood (Scots pine or Black pine), and called torch wood (Ast, 2001), therefore this interpretation fits with the archaeological evidence from the Bàsura Cave.

After a walk of approximately 150 m from the original opening of the cave and a climb of about 12 m, the group arrives at the *‘Corridoio delle impronte’*. They proceeded roughly in a row, the smallest left behind, and walked very close to the side wall of the cave, a safer approach also used by other animals (e.g. Canidae *incertae sedis* and bears) when in progress within a poorly lit and unknown environment. The slope of the tunnel floor, inclined about 24°, may have further forced the individuals to proceed in the only flat area, a couple of meters near the left wall of the cave, in the lower corridor. After about ten meters from the beginning of the *‘Corridoio delle impronte’*, the cave vault progressively drops below 80 centimeters and the members of the group were forced to crawl (Figure 9B), placing their hands (Figure 10) and knees (Figure 7b) on the clay substrate (Figure 8) (see also Movie, Appendix 2).

After some few meters, the group leader stopped, impressing two parallel calcigrade footprints, probably to choose the next movement and proceed forward to cross the lowest sector of the cavity vault. The other individuals stopped exactly when the leader stopped before, and then they proceeded along the same path by crawling following the group leader, as highlighted by the timing reconstructed from tracks interferences (Figure 8d3).

After passing a bottleneck of blocks and stalagmites, they descended for about ten meters along a steeply sloping surface. The whole group traversed a small pond, leaving deep tracks on plastic waterlogged substrate, climbed a slope of 10 meters beyond the *‘Cimitero degli orsi’*, and finally arrived at the terminal room *‘Sala dei misteri’*, where they stopped. On the walls, several charcoal traces coming from the torches are preserved.

Some charcoal handprints produced with a flexed arm placed more than 170 cm in height on the vault of the *‘Sala dei Misteri’* confirm that even the tallest individuals (Morphs. 4 and 5) reached this part of the gallery. The fact that their footprints are not preserved is related to the loss of the central portion of the hall floor. In the same room, the adolescent and the children started collecting clay from the floor and smeared it on a stalagmite at different levels according to their stature, as suggested by the breadth and relative distribution of the finger flutings on the karst structure. During the stay in the innermost room of the cave, the young individual that produced Morph. 2 imprinted ten clear heel traces (Citton et al., 2017), which are here interpreted as calcigrade tracks produced by a momentarily standing-still trackmaker involved in social activities comprising clay excavation and manipulation, as similarly mentioned for the ‘Salle des Talons’ at Tuc d’Audoubert cave (Pastoors et al., 2015).

After stopping for several minutes (considering the quantity and ubiquity of the tracks), they made an exit, following a route only partially adhering to the entry route. After passing the small pond, they crossed the upper corridor in a more comfortable and safe manner (Figure 9C). In fact, in the upper corridor all the prints show direction facing the exit while (Figure 11), with the axis of the foot parallel to the walls, as in the lower corridor most of the footprints are directed towards the interior of the cave.

### Concluding remarks

A holistic analysis of multiple and inter-related ichnological evidences allowed us to reconstruct several snapshots depicting a small and heterogeneous group of Upper Palaeolithic people that about 14,000 years ago explored a cave (Movie, Appendix 2). They overcame its uneven topography and carried out social activities in the most remote room, leaving evidence in their traces of a unique testimony to human curiosity.

The lower corridor was travelled in entry direction and documents the first unequivocal evidence of crawling locomotion in the human ichnological record, adopted by the explorers as a locomotor behaviour to obviate variation in the vault height from the trampled floor. Most probably, the group, chose to earn the exit from the cave through the upper corridor, easier for the higher vault height and the firmer substrate since no outgoing footprints are documented in the lower corridor. Obviously, in addition to this we must consider the exploratory factor, which has probably pushed the group forward simple for curiosity, along a different and still unexplored path to reach the cave exit.

Anatomical parts clearly registered on the substrate indicated that the individuals proceeded with naked limbs and that the body sizes were slender and muscular. Our study also confirms that very young children were active members during the activities of the Upper Palaeolithic populations, even in seemingly dangerous tasks, such as the deep exploration of the cave environment lit only with torches. As recently suggested for other European caves (Pastoors et al., 2015, 2017; Ledoux et al., 2017) the ‘Grotta della Bàsura’ site strongly supports the hypothesis that the cave exploration in Upper Paleolithic was carried out by heterogeneous groups.

The necropolis of the Arene Candide Cave (AMS dates spanning 12,820-12,420 cal BP for the first phase and 12,030–11,180 cal BP for the second phase), consisting of a “mixed” sample (males, females, adults, children), suggests an Upper Palaeolithic people composition very similar to those highlighted in the ‘Grotta della Bàsura’ (Sparacello et al., 2018). The burial of a newborn recently discovered in the Arma di Veirana cave (Erli, Savona, Liguria), in a valley 10 km from the coast, it still seems to emphasize those women and children systematically followed the movements of the group in the territory (F. N. pers. obs.) and shared, at least in part, the activities of men. The tracks left in the Bàsura indeed show us that hunters-gatherers behavior was not always driven by subsistence necessity, but as many ethnographic examples teach us, also by fun and ludic activities.

## Material and Methods

### Chronology

The first radiometric dating of charcoals fixed the human presence in the cave to the Upper Palaeolithic, around 12,340±160 years BP (De Lumley and Giacobini, 1985). The stalagmite crust preserving the footprints and incorporating fragments of coal is dated between 14,300±800 and 13,100±500 (Yokoyama, 1985). The last phase of the stalagmite growth, closing the entrance and sealing the ‘time capsule’, occurred at 12,000±1100 (Yokoyama, 1985).

New radiometric dating were performed in 2017 with the MICADAS AMS facility at Groningen (NL) on-charcoal samples of *Pinus sylvestris* (Table 1). Material was collected from the trampling palaeosurface during recent excavations inside the Mysterys’ Room.

### Laser Scanning acquisition

The whole record was contextualized into the rough topography of the environment and visualized through a three-dimensional mapping of the cave performed by laser-scanning. The main sectors of the cavity were digitally acquired via laser scanner ScanStation2 Leica and, more recently, ScanStation C10 Leica. The scans were performed at 360° (acquisition grid of the point cloud of 2x2 cm a probe 7 m and in correspondence to the areas with the highest concentration of traces, an acquisition grid of 0.5x0.5 cm probe 7 m). In total, 23 stations were run (9 in the *‘Sala dei misteri’* and 14 in the *‘Corridoio delle impronte’* areas); 38 targets (16 in the *‘Sala dei misteri’* and 22 in the *‘Corridoio delle impronte’*) were used for the point clouds registration. The Leica Geosystems HDS Cyclone 9.1 software was used to process the data. The recording shows a final alignment error of 2 mm for the model of the *‘Sala dei misteri’* and 1 mm for the *‘Corridoio delle impronte’* (Movie, Appendix 2). From the models, reliefs were obtained at various degrees of detail that allowed georeferencing all traces. The original cast performed in 1950 were digitally acquired via HDI Advance structured-light 3D Scanner R3x, with a resolution of 0.25 mm at 600 mm FOV (field of view). The data were processed with FlexScan3D Software (Figures 5, 6, 7d).

**Fig. 6.**
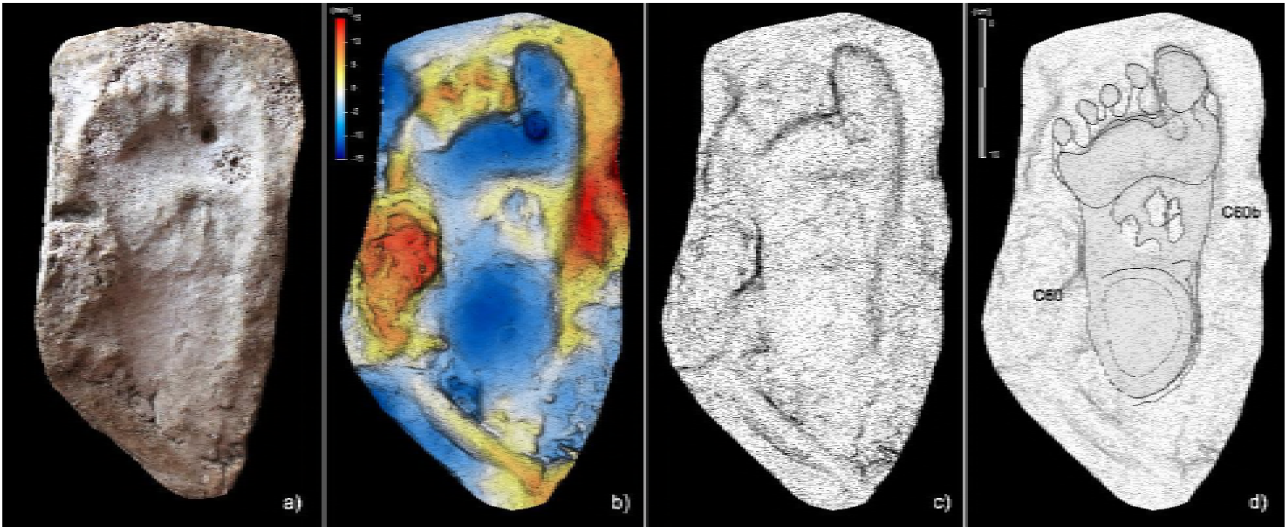
Plantigrade track from the ‘lower corridor’. **a**, cast of the 1950s reproducing the track C60 preserved in the sector A of the ‘lower corridor’ (see Fig. 1 main text). **b**, Digital terrain model of the cast obtained from the HDI 3D Scanner. **c**, Topographic profile with contour lines, obtained from b. **d**, Interpretive draw. A superimposed partial canid track, C60b, is clearly recognizable in the metatarsal area of the human footprint (Morph. 5).

**Fig. 7.**
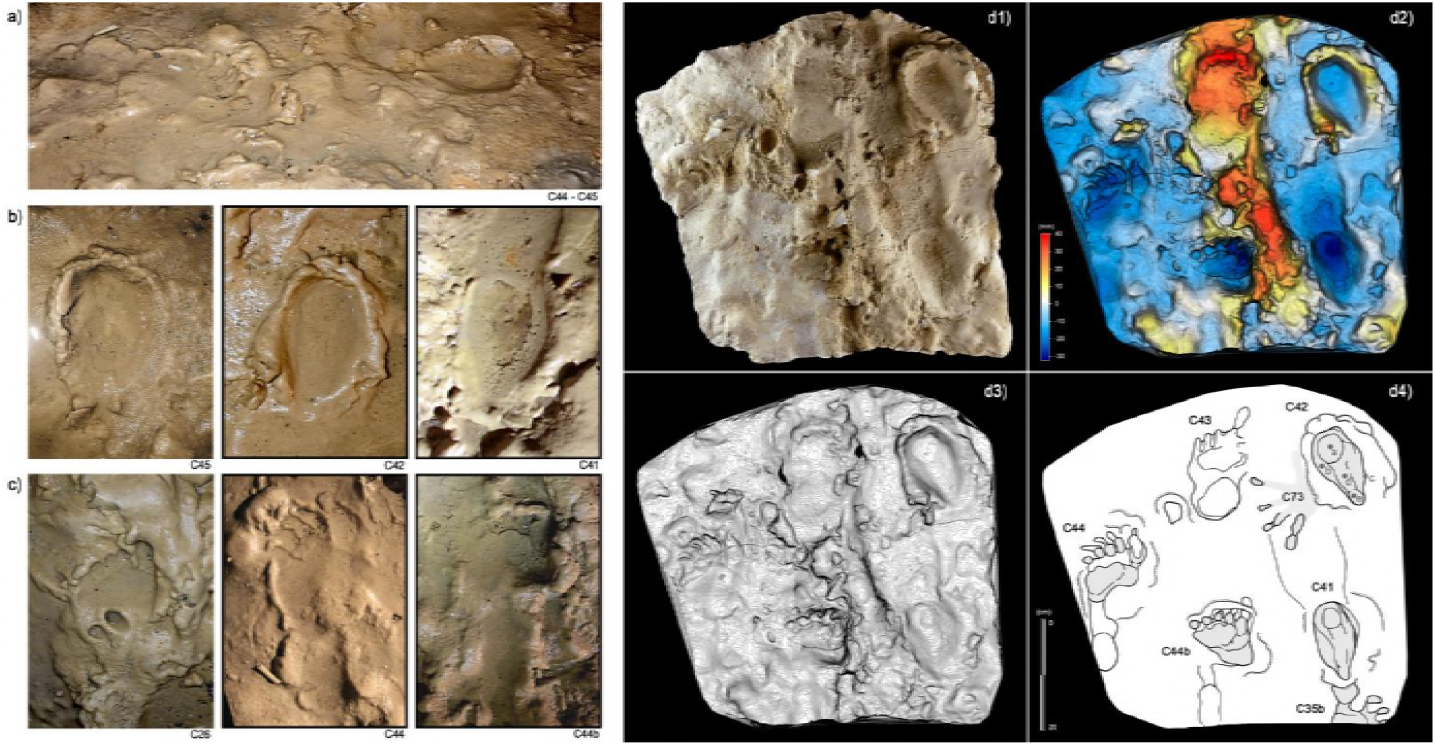
Selection of semi-plantigrade and knee traces from the ‘lower corridor’ of the *‘Corridoio delle impronte’* in the Bàsura cave, indicating crawling locomotion of the producers. **a**, Associated metatarsal (C44) and knee (C45) traces allowing estimation of the tibial length of the producer. **b**, Knee traces (C45, C42 and C41) imprinted on a plastic, waterlogged muddy substrate. **c**, Metatarsal traces (C26, C44 and C44b) imprinted on a plastic, waterlogged muddy substrate. **d1**, cast of the 1950s reproducing two knee (C41 and C42) and two metatarsal (C44, C44b) traces preserved in the area b of the ‘lower corridor’ (see Fig. 1). **d2**, Digital Terrain Model obtained from the HDI 3D Scanner. **d3**, Topographic profile with contour lines, obtained from d2. **d4**, Interpretive draw. In the knee trace C42 are located the impressions of the patella (a), vistas medals (b), the fibular head (c), the patellar ligament (d) and the tibial tuberosity (e).

### Digital Photogrammetry

Several photogrammetric models were subsequently obtained using several photos taken with 24 Megapixel Canon EOS 750D (18 mm focal length). The software used to build models is Agisoft PhotoScan Pro, (www.agisoft.com). High resolution Digital Photogrammetry is based on Structure from Motion (SfM) (Ullman, 1979) and Multi View Stereo (MVS) (Seitz et al., 2006) algorithms and produces high quality dense point clouds. The accuracy of the obtained models is up to 1 mm for close-range photography. The reconstructed 3D surfaces were then processed in the open-source software Paraview. False coloured models with contour lines, highlighting general morphology and differential depth of impression of the traces, were obtained (Figures 8a,b, 10, 11).

**Fig. 8.**
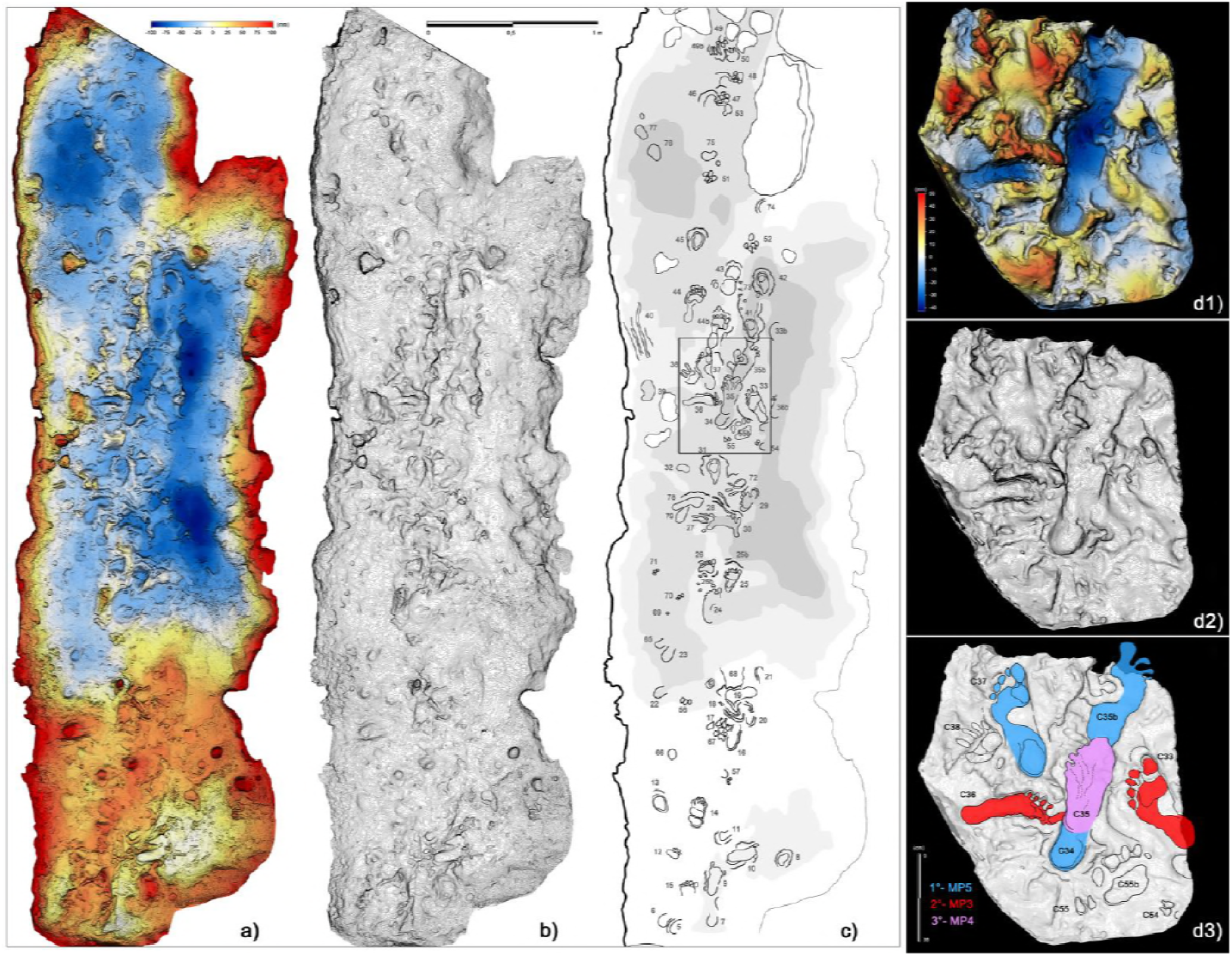
Crawling locomotion in the ‘lower corridor’ (sector B in Fig. 1). **a**, Colour topographic profile obtained from the digital photogrammetric model. **b**, Topographic contoured profile. **c**, Interpretive draw of the track-bearing surface (numbers identify single tracks and traces and are to be intended as preceded by the letter C). **d1**, Digital Terrain Model obtained from a cast of the 1950s reproducing a small area of the ‘lower corridor’. **d2**, Topographic profile with contour lines, obtained from d1. **d3**, Interpretive draw and timing of the different recognized tracks.

**Fig. 9.**
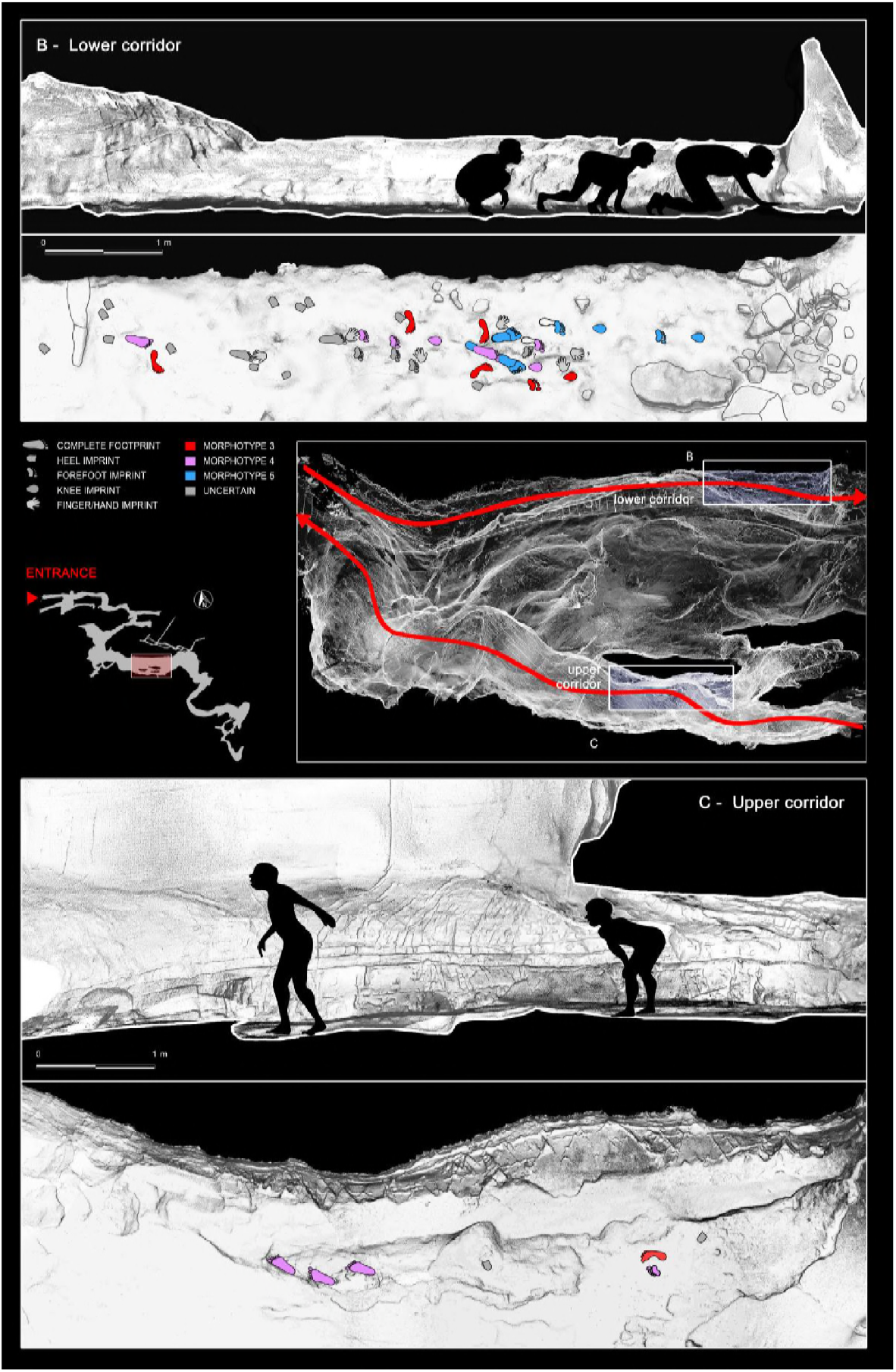
Reconstruction of the exploration routes chosen by the producers to enter and exit the cave. **B**, Crawling locomotion adopted by the producers to cross the ‘lower corridor’ and access to the innermost rooms of the cave. **C**, Exit route passing through the ‘upper corridor’, traveled by the producers in complete erect walking. The smallest producers are not reported in the sketch.

**Fig. 10.**
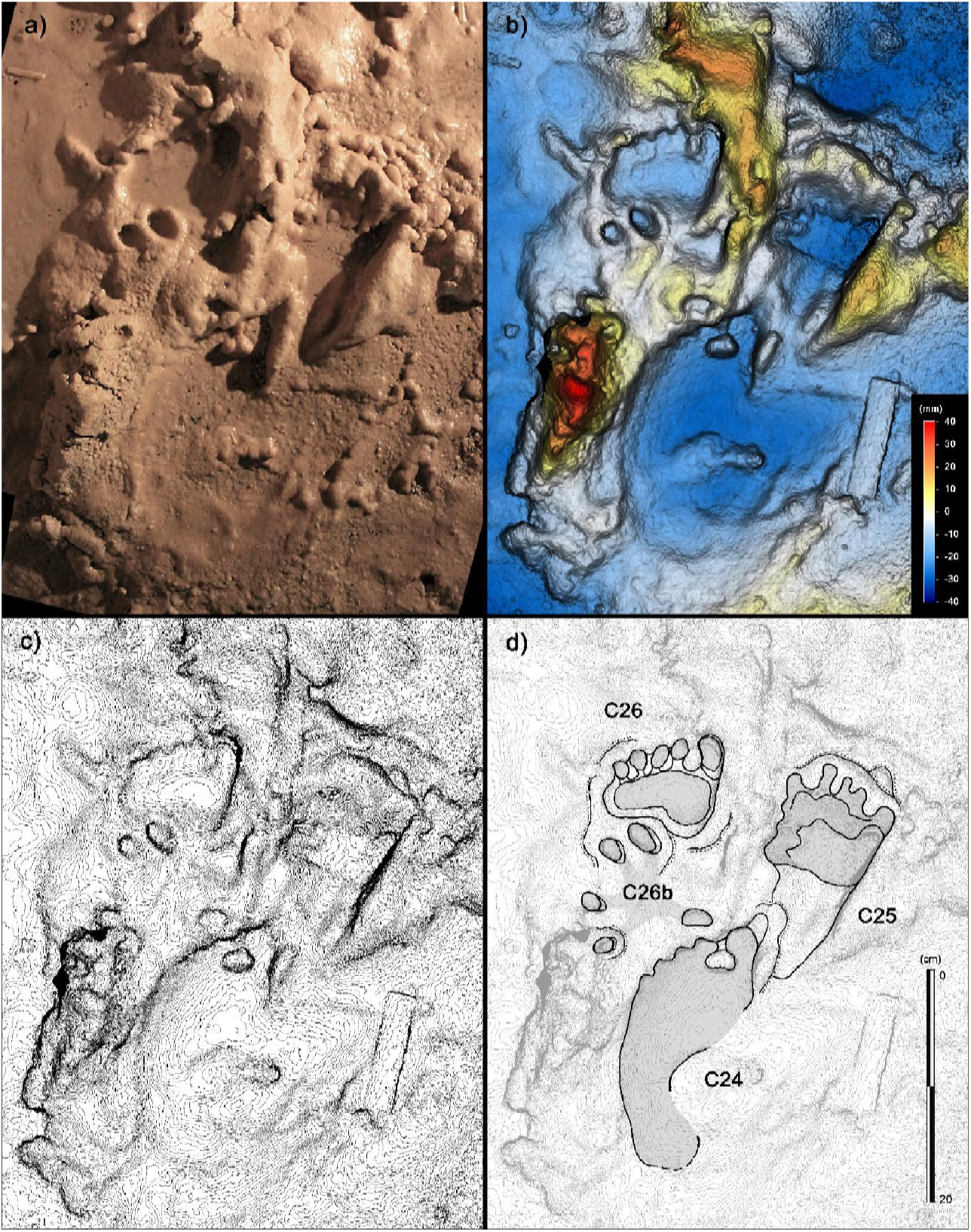
Human tracks from the ‘lower corridor’. **a**, Tracks C26, C26b, C25 and C24 from the sector B of the ‘lower corridor’ (see Fig. 1 main text). **b**, Digital terrain model obtained from high-resolution photogrammetry. **c**, Topographic profile with contour lines, obtained from b. **d**, Interpretive draw. C26b is interpreted as a partial hand-print of which only digit traces are preserved, interfering with a metatarsal trace deeply imprinted on a muddy, highly plastic, substrate.

**Fig. 11.**
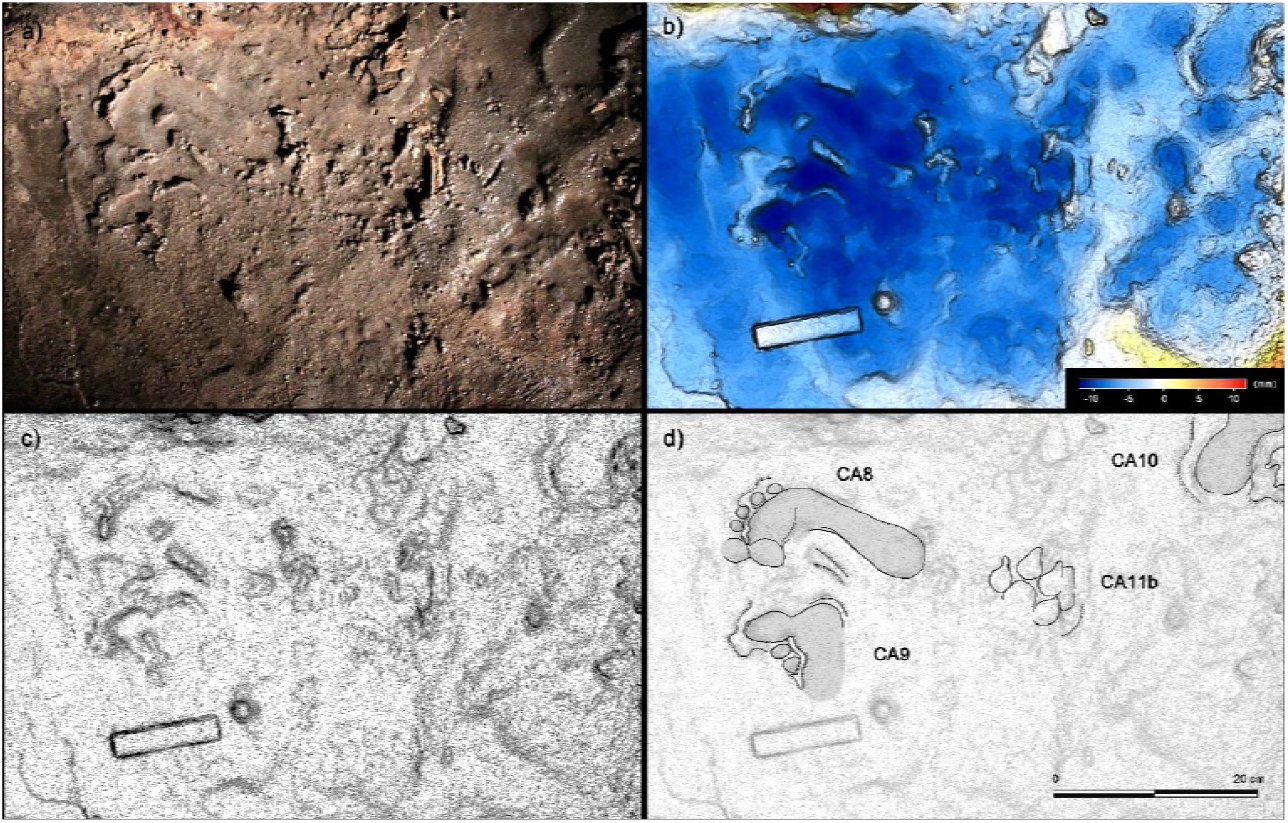
Shallow human tracks from the ‘upper corridor’. **a**, Tracks CA8, CA9, CA10 and Ca11b from the sector C of the ‘upper corridor’ (see Fig. 1 main text). **b**, Digital terrain model obtained from high-resolution photogrammetry. **c**, Topographic profile with contour lines, obtained from b. **d**, Interpretive draw. Tracks were impressed on a hard carbonate substrate covered by a thin muddy deposit, few millimeters in thickness.

### Analysis of human footprints

All recognized tracks (107 human traces) were analyzed directly on the field through a purely morphological approach using available landmarks (Robbins, 1985). The differential depth of each individual impression was analyzed directly in the field to infer a complex and multiphase biomechanics. Each isolated footprints, and those associated in trackways, were also drawn in the field on plastic film. The morphological and dimensional parameters directly collected were doublechecked, using photos and photogrammetric models. In addition the following two indexes were considered: Footprint index (FI), equal to foot width/foot length x100 and Arch angle (Aa), represented by the angle between the footprint medial border line and the line that connect the most medial point of the footprint metatarsal region and the apex of the concavity of the arch of the footprint (Clarke, 1933). In the reconstruction of body dimensions and age, only the foot measurements derived from the better-preserved footprints were used (Table 3).

### Principal Component Analysis

The 23 better preserved footprints, have been subjected to a Principal Component Analysis (PCA). For the analysis the software PAST 3.10 (Hammer et al., 2001) was used. Different homologous points were selected for the measures (Robbins, 1985) (Figure 12), namely nine anatomical lengths and widths (foot lengths (Dt1-BL, Dt2-BL, Dt3-BL); ball medial length (mtm-BL); ball lateral length (mtl-BL); heel medial length (ccm-BL); heel lateral length (ctul-BL); widths of ball (mtm-horiz.) and heel (ctul-horiz.) (Table 4). The raw data were log-transformed before the analysis to fit linear models and for the correspondence of the log transform to an isometric null hypothesis (Chinnery, 2004; Cheng et al., 2009; Romano and Citton, 2015, 2016; Romano, 2017). Missing entries were treated according to the ‘iterative imputation’ in PAST 3.10, preferable to the simple ‘mean value imputation’ (Hammer, 2013). The results of the PCA are reported in the scatter plots of Figure 4a whereas the loadings for the first three principal components are provided as supplementary information (and Appendix 1 Tables A, B).

**Fig. 12.**
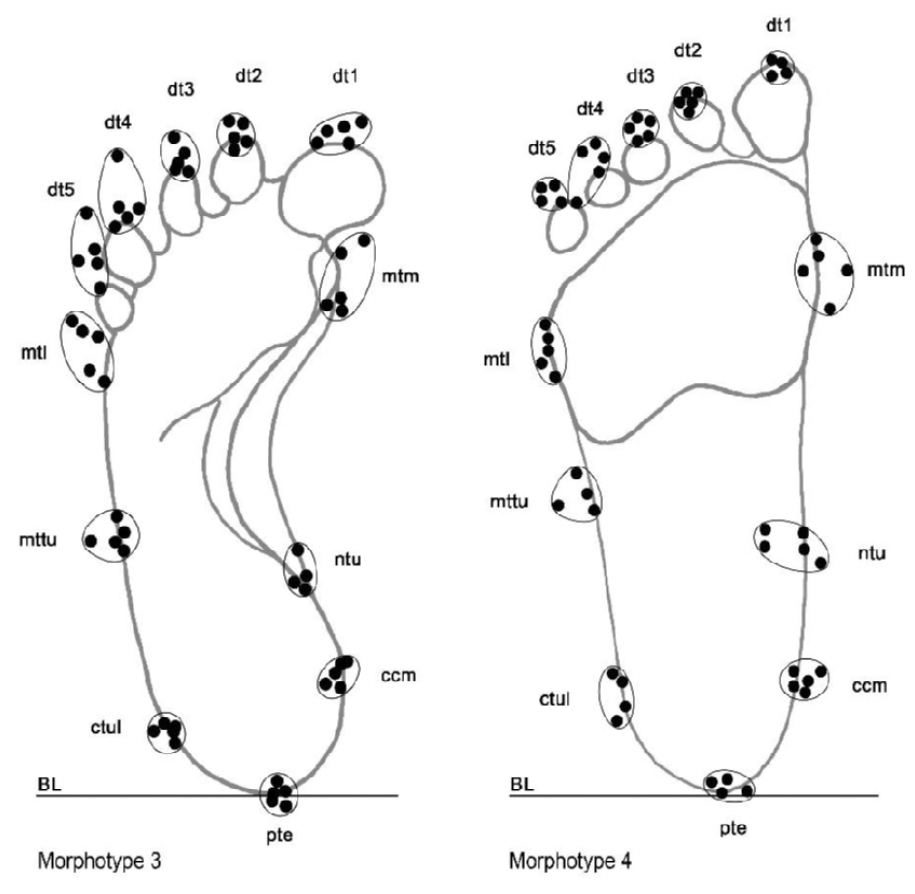
Adopted landmarks utilized to perform morphometric analysis, showed in two distinct mor-photypes (Morphs 3 and 4) as example. Landmarks in the distal portion of digit traces 4, 5, and in the medial, central and lateral portions of the sole trace were not considered reliable enough for the large variability, higher than the fixed error value (± 0.5 cm).

**Table 4.** Footprints and relative measures used for the Principal Component Analysis. Anatomical abbreviations as in Methods Section.

| ID | LENGTHS |  |  |  |  |  |  | WIDTHS |  |
| --- | --- | --- | --- | --- | --- | --- | --- | --- | --- |
|  | Dt1-BL | Dt2-BL | Dt3-BL | Ball medial<br>(mtm-BL) | Ball lateral<br>(mtl-BL) | Heel medial<br>(ccm-BL) | Heel lateral<br>(ctul-BL) | Ball<br>(mtm-horiz) | Heel<br>(ctul-horiz) |
|  | (cm) | (cm) | (cm) | (cm) | (cm) | (cm) | (cm) | (cm) | (cm) |
| SM3 | 13 | 12,3 |  | 10,5 | 9,2 | 2,3 | 2,4 | 5,5 | 4,3 |
| SM4 | 13,5 | 12,5 | 11,5 | 10,2 | 8 | 2,2 | 1,6 | 5,5 | 4,2 |
| SM43 | 13,5 | 13,5 |  | 10,5 | 9,4 | 2,2 | 1,8 | 6 | 4,6 |
| SM17 | 14,2 | 13,8 | 13,5 | 10,2 | 8,5 | 2 | 1,8 | 6,5 | 4 |
| SM5 | 17 | 16,8 | 15,5 | 12 | 12 | 2,5 | 2 | 6,5 | 5 |
| SM42 | 17 | 16,8 |  | 12,5 | 11,5 | 3,4 | 2,5 | 7 | 5,4 |
| SM26 | 18 | 16,5 | 15 |  |  | 2 | 3 |  | 5 |
| CA8 | 20,2 | 19,8 | 19,4 | 15 | 13 | 2,6 | 2 | 7,5 | 4,5 |
| C10 | 20,5 |  | 19,2 | 15,5 | 12,5 | 2,8 | 2,6 | 7 | 5,6 |
| SM15 | 20,5 | 20,5 | 18,5 | 15,4 | 13 | 4 | 2 | 7 | 6 |
| SM11 | 21 | 20,2 | 19,7 | 15,4 | 14,5 | 3,3 | 2,3 | 7,5 | 6 |
| SM6 | 21 | 21,5 | 20,5 | 17,5 | 15,5 | 4 | 2,2 | 9,2 | 5,7 |
| SM1 | 21,2 | 21,3 | 20,8 | 17,4 | 14,8 | 3,8 | 2,5 | 8 | 6 |
| C33 | 22,2 | 21 | 19,5 | 15,5 | 10,7 | 2,5 | 2 | 10 | 5,8 |
| C36 | 22,7 | 21,7 | 19,5 |  | 14,5 | 3 | 2,5 |  | 5,5 |
| M21 |  | 21,6 | 21,3 | 16,7 | 14,6 | 3,5 | 2,4 | 9,6 | 6,2 |
| CA1 | 22,4 | 21,8 | 20,8 | 14,8 | 12,8 | 3,1 |  | 8 |  |
| CA2 | 22,5 | 22 | 21 | 17,3 | 15,2 | 3,5 | 3,5 | 8,8 | 6,4 |
| C61 | 23 | 21,7 | 20,8 | 16,4 | 14 | 3,8 | 3 | 7,5 | 5,4 |
| C63 | 23,3 | 22,2 | 20,8 | 17 | 14 | 3,5 | 3,8 | 8,5 | 5,3 |
| C60 | 25,3 | 24,2 | 22,7 | 18 | 14,8 | 3,5 | 3,3 | 10,5 | 6,4 |
| C37 | 25,7 | 23,8 | 22,5 | 18,4 | 14,7 | 3,7 | 3,5 | 10,5 | 7 |
| C35B | 26,2 | 24,8 | 22,8 | 17 | 13,7 | 3,7 | 2,7 | 9,5 | 6,7 |

### Stature

Stature can be estimated from foot length (Robbins, 1985; Oberoi, 2006; Krishan and Sharma, 2007; Kanchan et al., 2008; Pawar and Pawar, 2012). Stature varies with race, age, sex, heredity, climate and nutritional status. Based on skeletal evidence it is thought that the body proportions of terminal Upper Palaeolithic individuals was similar to modern humans (Trinkaus, 1997; Ruff et al., 2005; Shackelford, 2007) but the foot length/stature ratio was regarded highly uncertain, between 0.15 and 0.16. Consequently, we calculated the foot length/stature ratio basing on a sample of terminal Upper Palaeolithic adult individuals (n.8) of Italian Peninsula (Corrain, 1977; Paoli et al., 1980; Formicola et al., 1990; Mallegni and Fabbri, 1995; Mallegni et al., 2000). The calculated ratio is found to be 0.1541, appearing very close to those proposed for modern humans between the XIX and XX centuries (Robbins, 1985; Topinard, 1978). Stature estimation from long bones length is commonly used in forensics medicine. In this study, we have used for the checking of the stature of the Morph. 5 the percutaneous length of tibia as they are known to give strong correlation to body height. We have used the relation S = 101.85+1.81 x PCTL± 3.73 for male and S= 77.86 +2.36 x PCTL± 2.94 for female (where S = stature and PCTL = Percutaneous tibial length) (Lemtur et al., 2017). For a tibial length of 35 cm, the stature of morhopyte 5 result of 165.2 ± 3.73 cm, compatible with the stature assumed from foot length (166.99 ± 2.93 cm).

### Body mass

Body mass estimates were derived from footprints parameters, basing on the assumption that body proportions have been constant through time (Dingwall et al., 2013). Regression formulae obtained from current dataset are based on mature individuals ranging between 154 and 185 cm in stature (Weight Kg=4.71+(1.82xFL)) (Dingwall et al., 2013; Bavdekar et al., 2006; Ashton et al., 2014) or on children (Grivas et al., 2008) with an average height of 147,44 cm (weigth Kg=-71.142+(5.259xrigthFL). We have used these formulae for the individual taller than 147 cm (Table 3), stature: (b) (Bavdekar et al., 2006), (c) (Grivas et al., 2008). For the three smaller individuals we used a dataset based on extant Caucasian children between 6 and 11 years old (n. 7147) ranging in stature between 118.6 and 145.7 cm (Malina et al., 1973), in order to develop a mathematical relation between foot length and body mass for young individuals. The report is nonlinear and expressed by the formula mass = 2.2897 e^0.126FL^ (Citton et al., 2017).

### Age

The foot length varies by age and sex. Studies on the relationship size/age of the foot in current juvenile individuals (Fryar et al., 2012; Müller et al., 2012) have highlighted that one year old individuals have a foot length equal to 13.07 ± 1.59 cm, reaching 24.4 ± 2.96 cm at the age of 13 years. The age estimation based on growth curve built on extant population is very similar.

However, we must consider that the reference anthropometric data mainly refer to modern well-nourished populations, with body size most likely higher at the same age. Anthropometric studies suggest that the morphology of the foot changes and becomes more elongated when the arch stabilizes around the age of six (Müller et al., 2012). In extant human populations, from the fifth or sixth years of age, arch angles vary from 21° (3-4 years) to 43° (9-11 years) in young males, whereas they vary from 26° (3-4 years) to 47° (9-11 years) in young females (Forriol and Pascual, 1950). As a result, morphotype 2, 3 seem to be similar, likely suggesting a corresponding similarity of age between producers of the two morphotypes. Growth curve based on extant population with an average height similar of those of the Late Upper Paleolithic provided an estimate of the age of the trackmakers. For the MP5 the wide and stout morphology is here interpreted as due to a completely adult stage with the partial collapse of the plantar arch.

### Gender

Sex determination starting from foot has been proposed using Foot index and threshold values. However, such approach is not entirely accepted and some researchers objected that the threshold value could vary significantly between populations, thus making it very speculative a gender estimate applied merely to foot morphology (Walia et al., 2016). These variations could be due to fact that anatomic structures of foot shows ethnical and regional variations owing to climatic factors, physical activities, socio-economic status and nutritional conditions. Despite the method uncertain arch angle and footprint morphology suggests a possible male as trackmaker of the largest footprint group. For the morphotypes 1, 2, 3, 4 any inference of gender result possible.

## ACKNOWLEDGMENTS

This work was supported by the Soprintendenza Archeologia Belle Arti e Paesaggio per la Città Metropolitana di Genova e le province di Imperia, La Spezia e Savona, Genoa, Italy and the Municipality of Toirano. The funders had no role in study design, data collection and analysis, decision to publish, or preparation of the manuscript. Matthew Bennet provided critical comments to an advanced manuscript draft. Jonah N. Choiniere revised the English language and provided suggestions about the manuscript’s structure. Part of the research was funded by the National Geographic Early Career Grant to M.R. (EC-53477R-18) “*A multidisciplinary approach to a unique human ichnological record from the Grotta della Basura (Toirano, Savona Italy)*”.

## Appendix 1

**Table A.** Scores obtained from the Principal Component Analysis

| ID | PC 1 | PC 2 | PC 3 | PC 4 | PC 5 | PC 6 | PC 7 | PC 8 | PC 9 |
| --- | --- | --- | --- | --- | --- | --- | --- | --- | --- |
| SM3 | -0.39913 | 0.10426 | -0.064014 | 0.014386 | -0.025072 | 0.016054 | 0.016716 | 0.011934 | 0.0074328 |
| SM4 | -0.50068 | -0.042109 | -0.014907 | 0.032287 | 0.0032614 | -0.029028 | 0.019643 | -0.026998 | 0.0033618 |
| SM43 | -0.40104 | -0.019701 | -0.0054895 | 0.018735 | 0.0016977 | 0.027812 | -0.0033284 | 0.0083841 | -0.0075861 |
| SM17 | -0.42507 | -0.0052715 | 0.061746 | 0.0063252 | -0.038382 | -0.016919 | -0.0164 | 0.0059965 | -0.0012712 |
| SM5 | -0.1976 | -0.035523 | -0.0043672 | -0.03242 | 0.024328 | 0.032498 | -0.025789 | -0.011295 | -0.0026746 |
| SM42 | -0.086943 | 0.013761 | -0.08784 | 0.050378 | -0.009608 | -0.0043334 | -0.025838 | 0.015148 | 0.0033864 |
| SM26 | -0.20626 | 0.16623 | 0.063691 | -0.021075 | 0.045203 | 0.0078808 | -6,97E-02 | -0.013948 | -0.00040047 |
| CA8 | -0.051293 | -0.079409 | 0.054639 | -0.097538 | -0.026619 | -0.014175 | -0.002064 | 0.014186 | 0.00038865 |
| C10 | 0.0058167 | 0.019522 | 0.0058477 | -0.035311 | 0.03647 | -0.00018801 | 0.032056 | 0.019886 | 0.0034661 |
| SM15 | 0.040758 | -0.12904 | -0.095157 | 0.017228 | 0.043206 | -0.034662 | 0.00082729 | -0.005932 | -0.0085001 |
| SM11 | 0.06668 | -0.064129 | -0.032745 | -0.023447 | 0.034482 | 0.026778 | -0.0049565 | 0.0063519 | 0.0068743 |
| SM6 | 0.15624 | -0.12859 | -0.045021 | 0.0020611 | -0.050337 | 0.019214 | 0.001418 | -0.0080637 | -0.0071906 |
| SM1 | 0.14791 | -0.061674 | -0.059992 | -0.0097849 | 0.00034344 | 0.0088016 | 0.019386 | 0.010087 | -0.0078662 |
| C33 | 0.0099173 | -0.09178 | 0.16838 | 0.043167 | 0.0021726 | -0.017624 | 0.012848 | 0.010468 | -0.0049062 |
| C36 | 0.094929 | -0.029812 | 0.032258 | -0.047049 | 0.0025717 | 0.013716 | -0.009842 | -0.02199 | 0.0015418 |
| M21 | 0.16838 | -0.0796 | 0.021892 | 0.019189 | -0.011351 | 0.022302 | -0.0073946 | 0.0015577 | 0.014584 |
| CA1 | 0.16391 | 0.21771 | -0.0025937 | 0.0040729 | 0.0067048 | -0.016137 | -0.017977 | 0.012136 | -0.0084853 |
| CA2 | 0.22089 | 0.072731 | -0.022013 | 0.006969 | -0.0012392 | 0.036927 | 0.012619 | -0.0038898 | -0.0063643 |
| C61 | 0.15271 | 0.023445 | -0.066418 | -0.034022 | -0.0050802 | -0.045788 | -0.0011026 | -0.0017506 | 0.007746 |
| C63 | 0.19883 | 0.1176 | -0.020327 | -0.025171 | -0.043065 | -0.029182 | 0.0070871 | -0.013733 | -0.00144 |
| C60 | 0.28381 | 0.030453 | 0.049268 | 0.0231 | -0.021083 | 0.01058 | -0.0018458 | -0.0051978 | -0.00067736 |
| C37 | 0.3109 | 0.046052 | 0.029704 | 0.054401 | -0.0019844 | 0.016719 | 0.011316 | -0.0085163 | 0.0054307 |
| C35B | 0.24634 | -0.045122 | 0.033459 | 0.033518 | 0.033381 | -0.031244 | -0.01731 | 0.0051775 | 0.0031501 |

**Table B.** Loadings for each principal components. **a**, Dt1-BL; **b**, Dt2-BL; **c**, Dt3-BL; **d**, Ball medial (mtm-BL); **e**, Ball lateral (mtl-BL); **f**, Heel medial (ccm-BL); **g**, Heel lateral (ctul-BL); **h**, Ball (mtm-horiz); **i**, Heel (ctul-horiz). Anatomical abbreviations as in Methods section.

|  | PC 1 | PC 2 | PC 3 | PC 4 | PC 5 | PC 6 | PC 7 | PC 8 | PC 9 |
| --- | --- | --- | --- | --- | --- | --- | --- | --- | --- |
| **a** | 0.36632 | -0.031724 | 0.28459 | -0.15437 | 0.31322 | -0.38384 | -0.061146 | -0.3656 | 0.61412 |
| **b** | 0.36721 | -0.086634 | 0.19468 | -0.1893 | 0.19705 | -0.27853 | -0.3069 | -0.13269 | -0.74546 |
| **c** | 0.34834 | -0.062518 | 0.10263 | -0.17626 | -0.035551 | -0.093333 | -0.12422 | 0.88253 | 0.16995 |
| **d** | 0.34521 | -0.15 | 0.03307 | -0.16341 | -0.13051 | -0.019176 | 0.88858 | -0.036893 | -0.14898 |
| **e** | 0.33106 | -0.13532 | -0.24619 | -0.50577 | -0.065819 | 0.67412 | -0.2218 | -0.19439 | 0.099588 |
| **f** | 0.35243 | -0.24194 | -0.73379 | 0.3425 | -0.19673 | -0.31842 | -0.1176 | -0.070489 | 0.051056 |
| **g** | 0.32186 | 0.93175 | -0.1145 | 0.050472 | -0.09853 | 0.019641 | 0.025317 | -0.030012 | -0.030921 |
| **h** | 0.30501 | -0.13213 | 0.50725 | 0.46978 | -0.57082 | 0.21548 | -0.15222 | -0.12269 | 0.031883 |
| **i** | 0.245 | -0.051134 | -0.0026745 | 0.53499 | 0.68278 | 0.40417 | 0.10113 | 0.098963 | -0.039702 |

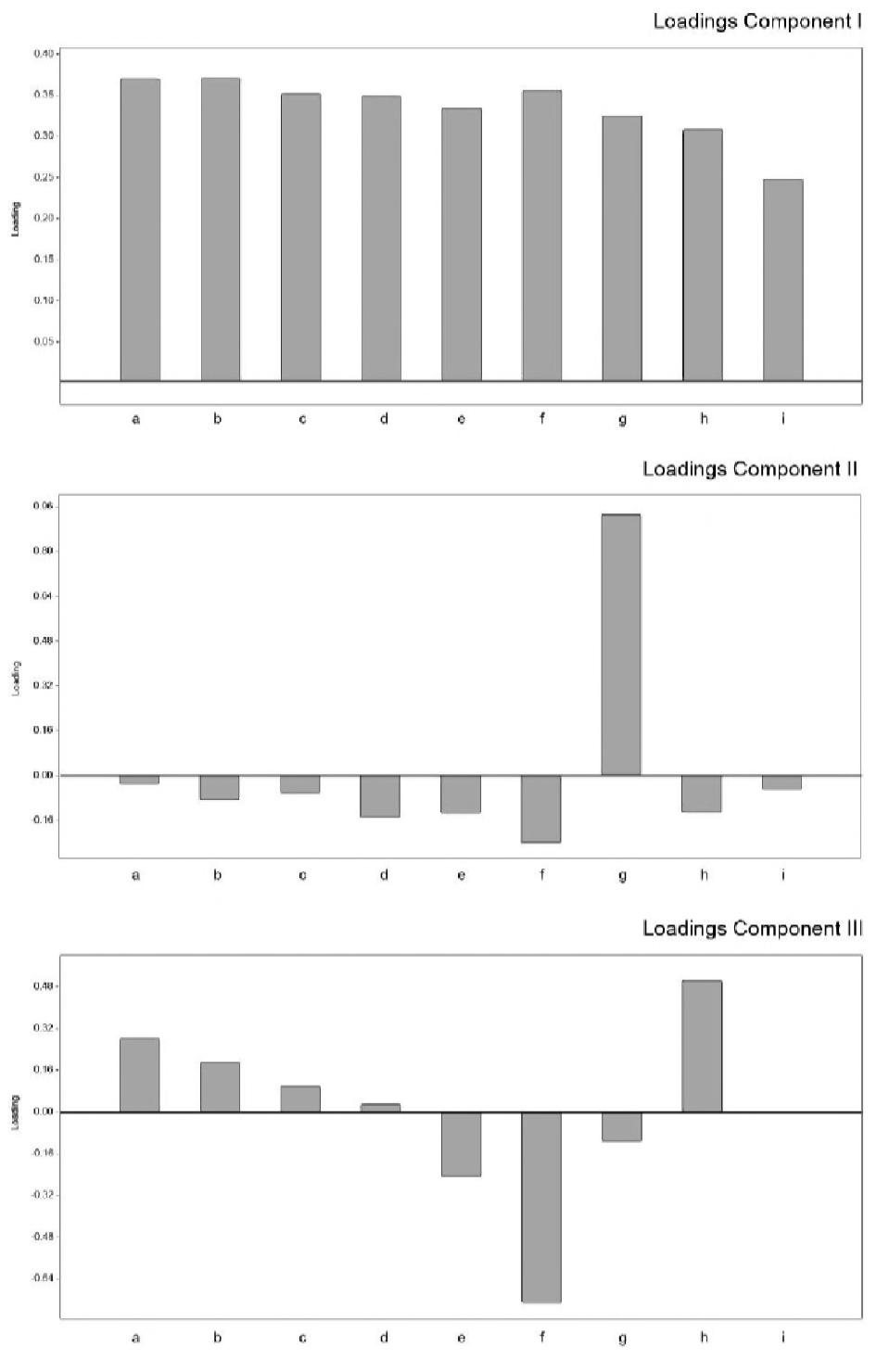
Loadings for the first three principal components. **a**, Dt1-BL; **b**, Dt2-BL; **c**, Dt3-BL; **d**, Ball medial (mtm-BL); **e**, Ball lateral (mtl-BL); **f**, Heel medial (ccm-BL); **g**, Heel lateral (ctul-BL); **h**, Ball (mtm-horiz); **i**, Heel (ctul-horiz). Anatomical abbreviations as in Methods section

